# Widespread collapse in Iberian forest site productivity projected under future climate change

**DOI:** 10.64898/2026.08.07.743474

**Authors:** Marta Fernández-Pastor, Gonzalo Rodríguez-Ruiz, Robert Monjo, María del Carre, Ana I. Hernández-Parada, Carlos Prado-López, Raúl García Valdés, Darío Redolat, Eulogio Chacón-Moreno, Jaime Ribalaygua

## Abstract

**Aim:** Here we aim to disentangle species-specific bioclimatic drivers of forest site productivity and project their future dynamics, providing a spatially explicit basis for anticipating climate-driven shifts in productivity and their implications for forest carbon sequestration.

**Location:** Iberian Peninsula.

**Time period:** 1985-2014 (calibration); 2071-2100 (projected under CMIP6 scenarios).

**Major taxa studied:** 21 Iberian tree species.

**Methods:** We used Site Form (SF) maps derived from the Third Spanish National Forest Inventory, spatially interpolating plot-level SF estimates as a continuous productivity index and relating them to 25 bioclimatic variables. Multiple linear regression models were selected via complementary stepwise and subset regression and validated on independent hold-out data (80%/20% split).

**Results:** Validated 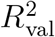 ranged from 0.46 (*Quercus faginea*) to 0.97 (*Pinus pinaster*); 17 of 21 species reached 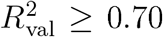. BI013 precipitation of the wettest month), not BI014, was the most frequently retained predictor (15/17); BI014 was retained in only (11/17 models with a near-even sign split. Combining projected changes in mean productivity and habitat extent under SSP5-8.5, fifteen of sixteen applicable species lose total productivity by 2071-2100, six — including *Fagus sylvatica* and *Betula alba* — collapsing to below 1% of their reference-period value; only *Pinus pinaster* gains, and only under the lowest-emission pathway (up to 175%) — under SSP5-8.5 it too loses productivity, albeit less than any other species (35% of its reference-period value retained). Limiting warming to SSP1-2.6 spares Mediterranean pine and oak species but not Euro-Siberian and montane ones.

**Main conclusions:** These validated, extrapolation-aware models reveal a near-universal, climate-driven collapse in Iberian forest site productivity, with direct implications for the carbon-sink potential currently attributed to these forest types, and provide a route to dynamic, climate-aware carbon-uptake estimates for the region.

## 1. Introduction

Forests cover approximately 30% of Earth’s land surface and store roughly half of terrestrial organic carbon, acting as a key buffer against anthropogenic CO_2_ emissions Pan et al., 2011). Yet this buffering capacity is not static. Forest carbon uptake depends on the balance between productivity and respiration, both of which respond sensitively to temperature, precipitation, and their seasonal distribution Baldocchi, 2008; Reichstein et al., 2013; Lindner et al., 2014). Under accelerating climate change, whether forests will remain net carbon sinks, become neutral, or shift to sources depends primarily on the bioclimatic control of growth (Hogan et al., 2024), making quantitative models of forest site productivity and their response to climate a prerequisite for credible carbon accounting.

The Iberian Peninsula sits at the intersection of Atlantic, Mediterranean, and continental climate regimes, sustaining some of Europe’s richest and most biodiverse forest ecosystems Hidalgo-Triana et al., 2023; Loidi, 2017). At the same time, it has been identified as one of the most sensitive regions to anthropogenic climate change, projected to experience warming above the European mean alongside a significant reduction in precipitation and intensification of summer droughts (Giorgi and Lionello, 2008; Lionello and Scarascia, 2018). A warming of 0.2-0.4 °C per decade has already been recorded since the 1970s across the Peninsula, accompanied by a lengthening of drought periods and an increasing frequency of extreme heat events (Pereira et al., 2021;, IPCC). These changes translate directly into ecological stress. Multispecies surveys across Iberian forests have documented widespread crown condition decline and amplified tree mortality from the late twentieth century to the present, strongly correlated with climate change-type drought, with more recent evidence confirming a continued increase in tree damage and mortality linked to drought intensity across Mediterranean forests of the Peninsula (Carnicer et al., 2011; Rebollo et al., 2024).

The relationship between climate and forest growth has been extensively studied through tree ring networks, which provide annually resolved, multi-century records of productivity. Across the temperate and boreal biomes, Babst et al. (2019) showed that the dominant climatic driver of tree growth was redistributed over the 20th century, with water availability replacing temperature as the primary constraint over large portions of the boreal zone and atmospheric water demand becoming increasingly limiting almost worldwide. For the Iberian Peninsula, which lies at the water-limited southern margin of this gradient, tree ring studies consistently show a dual response. Growth has increased in cold or high-elevation sites where temperature is the primary constraint, but has declined at warm, dry, low-elevation sites where drought stress dominates, particularly after the 1980s (Vilà-Cabrera et al., 2011).. Jump et al. (2006) documented rapid growth decline at the southern range edge of *Fagus sylvatica* in north-east Spain over the past decades, a pattern consistent with the climate-envelope contraction projected for this species under future warming. At a broader multispecies level, Ruiz-Benito et al. (2013) found that climate-driven mortality of Iberian trees was modulated by competition, with drought effects amplified in dense stands, confirming the interaction between bioclimatic conditions and stand dynamics.

Climate-oriented species distribution models (SDMs) have been widely applied to Iberian tree species to project future range shifts (García-Valdés et al., 2013; Benito Garzón et al., 2008, 2011), but they characterize where a species can occur rather than how productively it grows there, and therefore cannot capture the within-range gradients of site productivity that ultimately govern carbon uptake. Because a forest’s carbon sequestration potential is set by growth rates rather than by mere presence, resolving these gradients is essential for credible carbon accounting (Chisholm and Gray, 2024). Bridging this gap requires linking observed productivity data to bioclimatic gradients in a spatially explicit, species-specific way (Moreno-Fernández et al., 2018). Several studies have moved in this direction using National Forest Inventory NFI) data. For example, Charru et al. (2010) showed that long-term growth trends of common beech in France were positively associated with warming and rising CO_2_ but increasingly constrained by summer drought. In a complementary approach Seynave et al. (2005) modelled the site productivity of Norway spruce from environmental predictors, demonstrating that bioclimatic variables can explain a large fraction of landscape-scale site quality variation.

Forest site productivity is traditionally quantified through the Site Index (SI), defined as the dominant height at a reference age (Bontemps and Bouriaud, 2014). SI is a robust integrative measure of stand and growth, but it requires repeated measurements over long periods at permanent plots, limiting spatial coverage to a small fraction of the forest area (Calama et al., 2024). Alternative indices derived from static inventory records are therefore attractive for large-scale applications. Site Form SF), expressed as the dominant height of a stand at a reference dominant diameter, is an age-independent measure of site quality (Aguirre et al., 2022). SF maps at approximately 1 km resolution are available for 21 species (Aguirre et al., 2025), covering the commercially and ecologically most important forest types of the Iberian Peninsula and providing a continuous, interpolated dataset for bioclimatic modelling of site productivity.

European forests have seen their role as a continental carbon sink weaken in recent decades, with signs of saturation attributed to drought stress and disturbances outpacing CO_2_ fertilisation effects (Nabuurs et al., 2013). For the Iberian Peninsula, this signal is particularly concerning given the projected drying (Pereira et al., 2021; Gaitán et al., 2020). Current carbon sequestration calculators typically rely on static productivity factors per species and age class. One such example is the tool developed by the Spanish Office for Climate Change OECC) (Ministerio para la Transición Ecológica y el Reto Demográfico (MITECO), 2025). These static estimates cannot capture the expected changes in forest growth under future climates. Embedding species-specific bioclimatic regression models into such frameworks would transform static carbon inventories into dynamic projections sensitive to climate scenario, substantially increasing their relevance for climate policy and forest management planning.

Despite the availability of SF raster data for a broad set of Iberian species, no study has yet established species-specific bioclimatic regression models linking SF to climate across the peninsular Spain, evaluated their transferability through independent validation, or characterised the cross-species consistency of the controlling bioclimatic variables. These are the challenges we address here. Specifically, we relate observed SF rasters to 25 bioclimatic variables for the reference period 1985 – 2014 for the most commercially and ecologically important Iberian tree species. To this end we (i) fit and validate speciesspecific regression models using two complementary selection criteria stepwise AIC and subset regression BIC); (ii) quantify the selection frequency and sign consistency of each bioclimatic predictor to identify universal versus species-specific drivers; and (iii) apply the validated model coefficients to CMIP6 multi-model ensemble projections to map future changes in site productivity and their implications for forest carbon sequestration.

## 2. Material and methods

### 2.1. Site Form and bioclimatic data

Site Form rasters were collected from the Instituto Nacional de Investigación y Tecnología Agraria y Alimentaria (INIA-CSIC) for 21 tree species covering the peninsular Spain Fig. 1b; Table A.4) (Aguirre et al., 2025). These maps were derived from the Third Spanish National Forest Inventory (SNFI, 1997∓2007) and represent modelled SF values across the forest area occupied by each species, so that valid pixels are confined to the mapped distribution of the target species and coverage is therefore spatially discontinuous (Aguirre et al., 2022). Valid pixels were confined to each species mapped distribution (spatially discontinuous), at ~1 km resolution (WGS84), with total valid-pixel counts ranging from 908 (*Abies alba*) to 128 404 (*Quercus ilex*). Species exceeding 5000 pixels were randomly subsampled to *n* = 5,000 for model fitting, which helped keep computation tractable while maintaining distributional representativeness.

**Figure 1.**
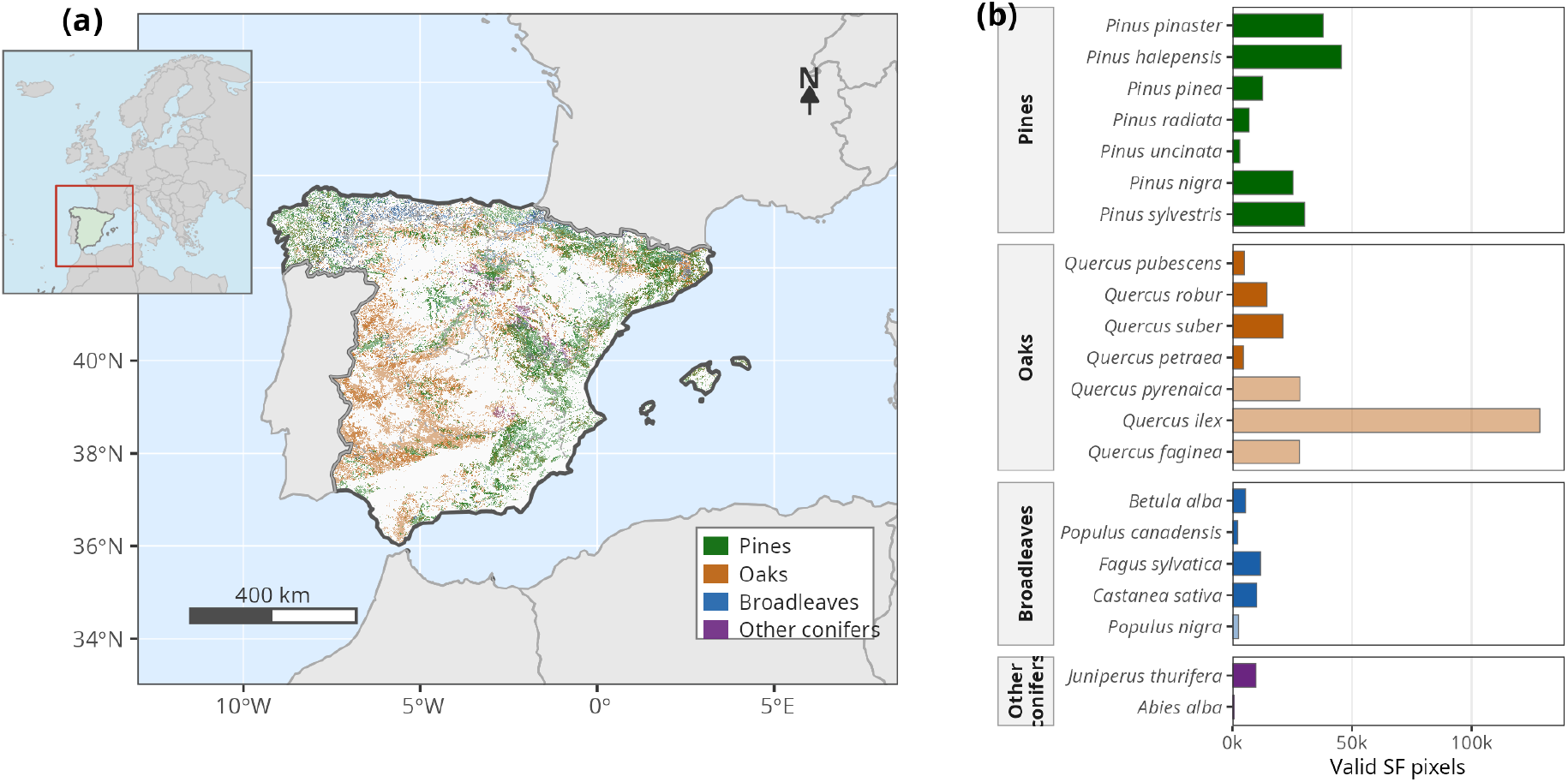
Study area and spatial coverage of Site Form (SF) data. (a) Geographic distribution of SF observations across the Iberian Peninsula derived from the Spanish National Forest Inventory (IFN). Each semi-transparent layer shows the spatial extent of SF data for one of the four taxonomic groups: pines (green), oaks (orange), broadleaved species (blue), and other conifers (purple); overlapping areas indicate co-occurrence of groups at the same ~ 1 km pixel. Dashed lines indicate autonomous community boundaries. Inset: location of the study area within Europe. (b) Number of valid SF pixels per species (total IFN count prior to subsampling), coloured by taxonomic group. Faded bars indicate species whose validated models did not meet the performance threshold applied for spatial projection (see Section 2.2).

Twenty-five bioclimatic variables for the reference period 1985–2014 were derived by a two-step analogue/regression statistical downscaling method (Ribalaygua et al., 2013) applied to high-resolution gridded climate data for the Iberian Peninsula: the 19 standard WorldClim-type variables (BIO01-BIO19, with dry periods fixed to physically consistent bounds to avoid unrealistic future values (see Gaitán and Pino-Otín, 2023; Chacón-112 Moreno et al., 2026)) and six evapotranspiration-related variables incorporating a topographic correction (aspect and slope) (BIO41-BIO46: annual, driest-month, wettest-month, warmest-month, coldest-month ET0, and ET0 seasonality). Bioclimatic values were extracted at each valid SF pixel by exact cell-centroid matching (`terra::extract`), yielding a per-species data frame with columns *{*SF; BIO01; …; BIO46*}*.

### 2.2. Model calibration and inference

#### Model calibration

For each species we fitted multiple linear regression models of the form

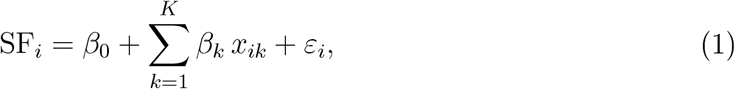

where *x*_*ik*_ are the selected bioclimatic predictors and *ε*_*i*_ ~ *N* (0, *σ*^2^). All predictors were retained on their original scale, without transformation.

Two complementary model-selection strategies were used. The first was bidirectional backward-forward) stepwise regression starting from the full model and retaining the predictors according to the Akaike s Information Criterion (AIC) `MASS::stepAIC`); this strategy prioritises predictive fit and typically retains a larger predictor set (Burnham and Anderson, 2002). The second was exhaustive best-subset selection which minimises the Bayesian Information Criterion (BIC), with the number of retained predictors constrained to a maximum of 11; this approach penalises complexity more strongly than AIC selection and favours parsimony for out-of-sample projection (Burnham and Anderson, 2002).

The data were split into training (80%) and hold-out (20%) sets using a stratified random partition based on quantiles of the response variable, so that the full range of SF values was represented in both sets. Predictive performance was assessed on the hold-out set using the coefficient of determination 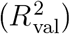 and root mean squared error SE):

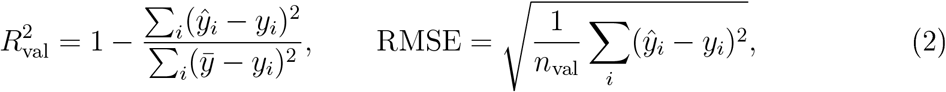

where *ŷ*_*i*_ are the model predictions and 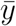 is the hold-out mean. Overfitting was diagnosed by comparing the training adjusted coefficient of determination 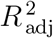

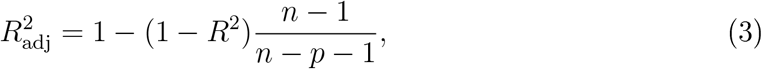

where *n* is the training sample size and *p* the number of retained predictors, against 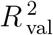. Models with 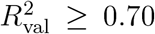 were considered sufficiently accurate for spatial application, consistent with validation thresholds reported in comparable site-productivity modelling studies (Bontemps and Bouriaud, 2014; Moreno-Fernández et al., 2018); species below this threshold are reported for completeness but excluded from spatial projection.

In addition, a third prediction approach was constructed for each species as the pixel-wise arithmetic mean of the AIC and BIC predictions at each pixel hereafter MEAN). This provides a simple model-ensemble estimate that can reduce individual model bias when the two selection strategies capture complementary aspects of the bioclimatic signal (Burnham and Anderson, 2002).

The three candidate predictions (AIC, BIC, and MEAN) were then retained for validation against SF data, and the approach with the lowest external RMSE was adopted as the final predictive model for each species, except where additional checks for spatial extrapolation or map degeneration justified an alternative choice.

#### Validation against SF data

As an independent test of predictive performance, the three candidate approaches (AIC, BIC, MEAN) were evaluated against the original SF dataset (Aguirre et al., 2022), which provides field-measured SF values at georeferenced plots not used during model calibration. This external validation covered 17 of the 21 species studied, the four species with 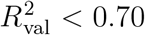 having been excluded from this step. For each species and each candidate approach, SF was predicted at the original plot locations using the reference period (1985-2014) climate bioclimatic layers, and the residual relative to the observed SF value was computed. The external RMSE was then calculated across all observed plots for that species, and the approach yielding the lowest external RMSE was adopted as the final predictive model and applied for all subsequent spatial products and climate projections (Table A.5).

This RMSE-based criterion was, however, overridden for two species, because both internal and external RMSE are evaluated only at known INIA plot locations and therefore cannot reveal whether a model extrapolates reliably once applied across a species full potential range. The limitation became apparent when constructing the final calibrated maps. *Juniperus thurifera*, despite an excellent statistical fit, extrapolated to unrealistic values across virtually its entire habitat-suitable area under reference-period climate and was therefore excluded from the spatial results reported below. *Pinus pinaster* showed a milder form of the same behaviour under its selected AIC model, so its reference-period and future projections instead use the more parsimonious BIC coefficients, which extrapolate far more conservatively while leaving mean productivity essentially unchanged. All analyses were performed in *R* 4.x (R Core Team, 2024) using the packages `terra`(Hijmans et al., 2026) for raster processing, `MASS`(Venables and Ripley, 2002) for stepwise AIC selection, leaps (Lumley and Miller, 2023) for best-subset BIC selection, and the tidyverse collection (Wickham et al., 2019) for data handling and visualisation.

### 2.3. Future climate projections

The regression coefficients of the validated models for each species were first applied to the reference-period bioclimatic layers to predict SF across the entire Iberian Peninsula and the Balearic Islands, rather than only at the calibration locations. The same coefficients were then applied to bioclimatic projections from ten Earth System Models ESM) comprising ACCESS-CM2, BCC-CSM2-MR, CanESM5, CMCC-ESM2, CNRM ESM2-1, EC-Earth3, MPI-ESM1-2-HR, MRI-ESM2-0, NorESM2-MM, and UKESM1-0LL (, IPCC). Projections were generated for four emission scenarios (SSP1-2.6, SSP2-4.5, SSP3-7.0, and SSP5-8.5) (O’Neill et al., 2017) and three future time horizons 2021–2050, 2041–2070, 2071–2100), yielding 120 scenario-period-ESM combinations per species. For the species whose final model was the MEAN approach, future projections were obtained by applying the AIC and BIC coefficients separately to each combination of ESM, SSP and time horizon, and then averaging the two resulting values at each pixel.

To synthesise the projections across ESMs for each scenario and time horizon, while accounting for the associated inter-model uncertainty, the median SF value across the ten ESMs was computed at each pixel. This produced a single median SF projection that is more robust to individual model biases than any single-model output (Knutti and Sedláček, 2013; Chacón-Moreno et al., 2026).

### 2.4. Percentage rescaling

The resulting baseline SF map for each species, described above, was rescaled and clipped as described below, and the resulting map was used as the final reference-period product. The same rescaling was subsequently applied to every future scenario, so that all products share a common scale.

All predicted SF rasters, including the reference-period prediction, the individual ESM projections, and the resulting median projections, were transformed onto a common scale. Because the native SF units differ markedly among species, the raw predictions are not directly comparable. For each species, we therefore expressed all predicted values on a common percentage scale defined by the range of SF observed in the INIA calibration data (Aguirre et al., 2022). The observed minimum was set to 0% and the observed maximum to 100%, and each predicted pixel value was rescaled linearly within this range. Predicted values falling below the observed minimum or above the observed maximum were then set to 0% and 100%, respectively, so that the final maps do not extrapolate beyond the range of SF represented in the calibration data. The same rescaling was applied identically to all rasters, so that every map shares a common scale.

### 2.5. Habitat suitability

For each of the 17 applicable species, a species distribution model (SDM) for the reference period was calibrated (REF). The presence data were obtained and curated from Third National Forest Inventory (IFN3) and the Spanish Forest Map (MFE50, 2015 version), and the distribution range was broader than the SF data plots. SDMs were calibrated and projected using the methods and variables described in Chacón-Moreno et al. (2026) for the Iberian Peninsula and the Balearic Islands. For some species for which presence data were available at the level of ecological genetic groups, which better reflect the local and regional adaptation of populations (Table 1), the SDMs were calibrated for these groups rather than for the species as a whole. For each taxon, the SDMs were projected onto future conditions for the same combination of ESM, SSP and time hori on, thereby producing suitability maps for the reference period and for the future presence/absence). These suitability maps were produced with the aim of restricting the maps inferred and projected on the basis of the shape index to those locations where the species is predicted to find suitable habitat, thereby reducing the potential extrapolation of the inferred shape index values.

**Table 1.** Species for which the SDM was calibrated at the level of ecological genetic groups rather than for the species as a whole, with the name of each modelled group. The remaining 8 applicable species (*Abies alba, Betula alba, Castanea sativa, Pinus radiata, Pinus uncinata, Populus canadensis, Quercus pubescens*, and *Juniperus thurifera*) were modelled as a single unit.

| Species | Genetic groups |
| --- | --- |
| *Pinus halepensis* | Central and southern Spain (gg1); Balearic Islands (gg2); southern Spain and Morocco (gg3); central and northern Spain (gg4); northern Spain and France (gg5) |
| *Quercus suber* | Northwestern (gg1); western core (gg2); southwestern core (gg3); southern Spain (gg4); eastern (gg5) |
| *Pinus pinaster* | Atlantic Iberian (gg1); central Spain (gg2); eastern Spain (gg3); southern Spain (gg4) |
| *Pinus pinea* | Northern Meseta (gg1); eastern (gg2); Andalusia (gg3); Catalonia (gg4) |
| *Fagus sylvatica* | Galicia (gg1); Pyrenees (gg2); central Spain (gg3) |
| *Pinus nigra* | Northeastern (gg1); central (gg2); southeastern (gg3) |
| *Pinus sylvestris* | Western (gg1); Pyrenean (gg2); eastern (gg3) |
| *Quercus petraea* | Western (gg1); Basque–Navarrese/interior ranges (gg2); eastern Pyrenean (gg3) |
| *Quercus robur* | Cantabrian (gg1); western Pyrenean/Iberian Range (gg2); eastern Pyrenean (gg3) |

Then the rescaled prediction was restricted to each species’ suitable area. Pixels where the SDM predicts absence are excluded regardless of the SF value the regression would otherwise return. This constrains every SF prediction to the multivariate climatic niche of the species rather than to the univariate range of individual bioclimatic predictors, and is applied identically to the reference-period baseline and to every future scenario.

Changes in SF between the current-climate baseline (1985–2014) and each future scenario are expressed both as absolute differences (SF units) and as percentile shifts within the species-specific INIA distribution, consistent with the presentation framework described in Section 4.4.

## 3. Results

### 3.1. Model performance and bioclimatic predictors

0f the 21 species modelled, 17 reached 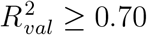 and were retained for spatial application (Table A.4); across these species, bioclimatic variables explained 78% of landscape-scale SF variance on average (range 0.44–0.97). External validation against independent INIA data selected AIC as the final model for 7 species, BIC for 6, and MEAN for 4 (Table A.5), reducing external RMSE by up to 59% relative to the best individual model where MEAN was selected. Full per-species regression coefficients are given in Supplementary Table S2.

Variable importance was assessed from each species’ actual final model (AIC, BIC, or the average of both for MEAN; Table A.4), rather than from AIC alone, so that the result reflects the models actually used for spatial projection; for MEAN species a variable counts as retained if selected by AIC, BIC, or both, with its sign taken from the average of the two coefficients (0 where a variable was not selected by one of them). Figure 2 shows, for each of the 23 bioclimatic variables retained in at least one final model, how many of the 17 applicable species’ models retain it (bar length) and the split between positive and negative coefficients among those retentions (bar colour).

**Figure 2.**
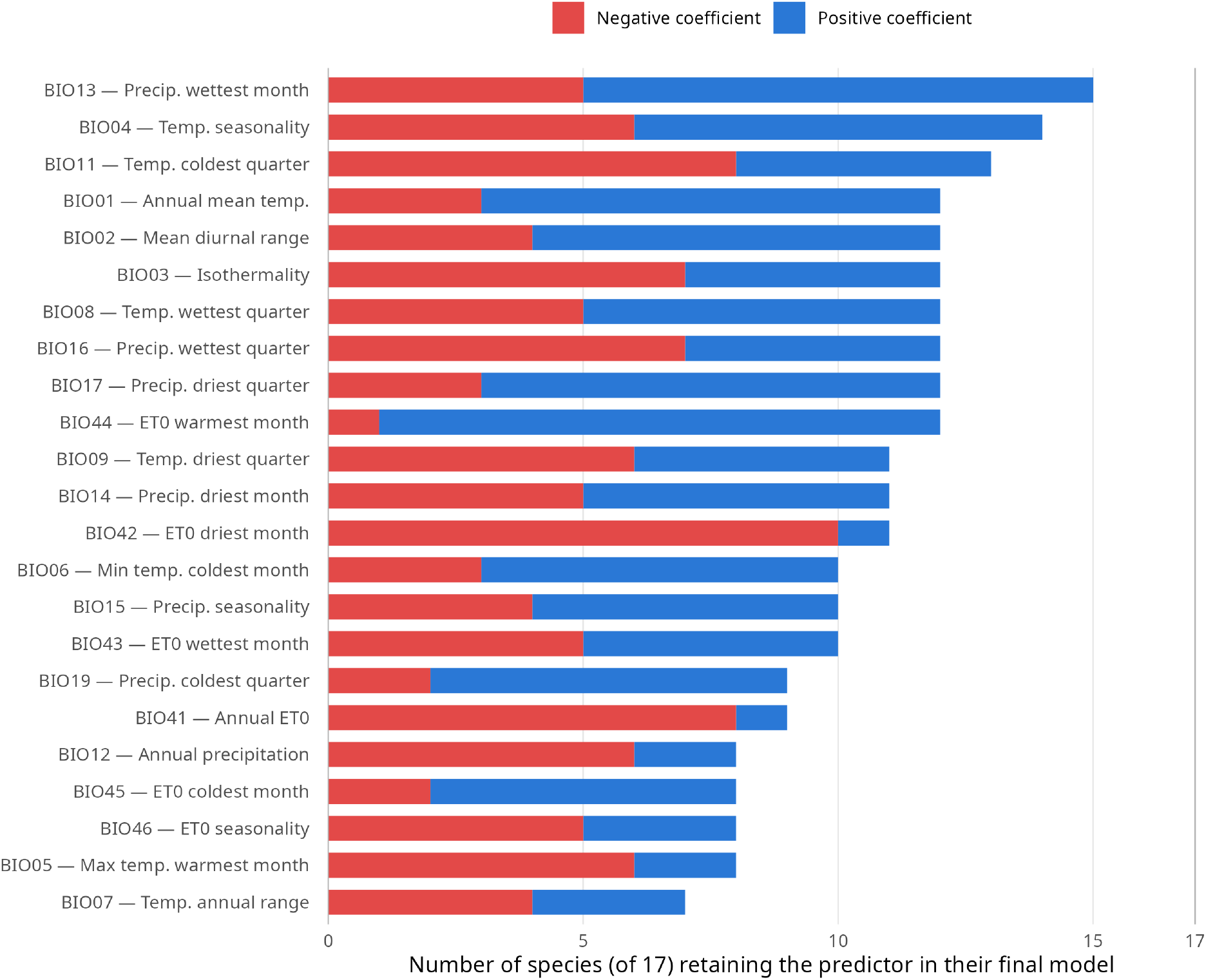
Bioclimatic predictor frequency and sign split across the 17 applicable species’ final selected models (AIC, BIC, or MEAN). Bar length is the number of species retaining each variable; colour indicates the split of species between positive (blue) versus negative (red) coefficient. BIO10 and BIO18 were not retained in any final model and are omitted.

BIO13 (precipitation of the wettest month) is the most frequently retained predictor (15/17), followed by BIO04 (temperature seasonality, 14/17) and BIO11 (temperature of the coldest quarter, 13/17, predominantly negative). Among predictors retained in at least 8 species, BIO44 (evapotranspiration of the warmest month) shows the strongest consistency of positive signs (92%), while BIO42 and BIO41 (evapotranspiration of the driest month and annual evapotranspiration) are predominantly negative (9% and 11% positive). BIO14 (precipitation of the driest month) — the flagship predictor when variable importance is assessed from AIC models alone — is retained in only 11 of 17 final models with a near-even sign split (55% positive), indicating genuine ecological ambivalence rather than a universal role once the actual production models are considered. BIO10 and BIO18 were not retained in any final model.

### 3.2. Spatial patterns in reference-period Site Form maps

Applying the calibrated regressions to the baseline of the reference-period reveals substantial variation in productivity across the 16 applicable species. Table 2 summari es these reference period (1985–2014) Site Form patterns, ranked by mean SF within each species’ own SF-mapped area. Productivity varies more than eight times between species, from 8.2% (*P. pinaster*)to 66.7% (*B. alba*). Three species – *B. alba* (66.7%), *A. alba* (64.7%) and *Q. pubescens* 63.8%) — form a distinct high-productivity group well above the rest; a broad middle tier of pines and *Quercus* species clusters between 44% and 53%; and a lower tier — *Q. suber, Q. robur, P. pinea, P. canadensis, P. pinaster*— falls between 8% and 28%.

**Table 2.** Reference period (1985–2014) Site Form patterns for the 16 applicable species, ranked by mean SF within the SF-mapped area. *r*_*lat*_/*r*_*lon*_: Pearson correlation between pixel SF and latitude/longitude within the species’ own range (only reported where |*r*| ≥ 0.2). *BIC-corrected model (see caveat above).

| Species | Suitable area (px) | Mean SF% | Dominant gradient within range |
| --- | --- | --- | --- |
| *Betula alba* | 59,943 | 66.7 | west ($r_{lon}=-0.61$) |
| *Abies alba* | 11,534 | 64.7 | west ($r_{lon}=-0.33$) |
| *Quercus pubescens* | 56,327 | 63.8 | south-east ($r_{lat}=-0.32$, $r_{lon}=+0.60$) |
| *Pinus nigra* | 74,309 | 53.4 | north-east ($r_{lat}=+0.37$, $r_{lon}=+0.54$) |
| *Pinus uncinata* | 19,801 | 53.1 | none |
| *Pinus radiata* | 41,542 | 52.6 | east ($r_{lon}=+0.48$) |
| *Pinus halepensis* | 154,834 | 50.1 | north-east ($r_{lat}=+0.38$, $r_{lon}=+0.52$) |
| *Quercus petraea* | 86,963 | 46.1 | north ($r_{lat}=+0.33$) |
| *Castanea sativa* | 81,530 | 45.5 | south-west ($r_{lat}=-0.23$, $r_{lon}=-0.22$) |
| *Pinus sylvestris* | 71,212 | 44.2 | north ($r_{lat}=+0.22$) |
| *Fagus sylvatica* | 65,399 | 43.9 | east ($r_{lon}=+0.55$) |
| *Quercus suber* | 141,089 | 27.8 | north-west ($r_{lat}=+0.81$, $r_{lon}=-0.23$) |
| *Quercus robur* | 76,275 | 25.9 | west ($r_{lon}=-0.48$) |
| *Pinus pinea* | 67,920 | 21.0 | south ($r_{lat}=-0.39$) |
| *Populus canadensis* | 71,761 | 16.3 | east ($r_{lon}=+0.52$) |
| *Pinus pinaster* * | 104,772 | 8.2 | north-west ($r_{lat}=+0.51$, $r_{lon}=-0.50$) |

Ten of the 16 species show a positive within-range correlation with latitude, i.e. higher SF toward the cooler, moister north of their Iberian distribution, including drought-tolerant Mediterranean species such as *P. halepensis* and *P. nigra* — consistent with the consistently positive sign of annual mean temperature (BIO01) discussed below being bounded by a thermal optimum rather than acting as an unconditional growth driver. The clearest case is *Q. suber* (r=0.81): cork oak is Mediterranean at the biome scale, but *within* its Iberian range the wetter, cooler northern populations clearly outproduce the hot, dry southern ones. A smaller group — *Q. pubescens, Castanea sativa, Pinus pinea* — instead shows higher SF toward the south of their (more restricted, higher-elevation or Atlantic-influenced) ranges. East–west gradients are similarly species-specific: Atlantic-affiliated species (*B. alba, A. alba, Q. robur, Q. suber* and the BIC-corrected *P. pinaster*) are more productive toward the humid west, while *Q. pubescens, Fagus sylvatica, Pinus nigra* and *Populus canadensis* are more productive toward the drier but more continental east. A within-range gradient this weak (below |*r*| = 0.2 in both directions) genuinely means no detectable directional trend in SF among a species’ own suitable pixels — it is not a statement about where the species occurs geographically, which for a narrow-range specialist like *P. uncinata* (confined to the Pyrenees) can still be highly localised.

### 3.3. Future changes: patterns of decline and resilience

Figure 4 tracks, for each species, a total productivity index — the product of mean SF and SF-mapped area, normalised to the reference-period baseline (set to 100%) — through the three future horizons, under each of the four emission scenarios (species ordered as in Table 2, from highest to lowest reference-period productivity). Because it combines both dimensions of change, this index captures productivity collapse that neither area nor mean SF alone would reveal (Section 4.3).

**Figure 3.**
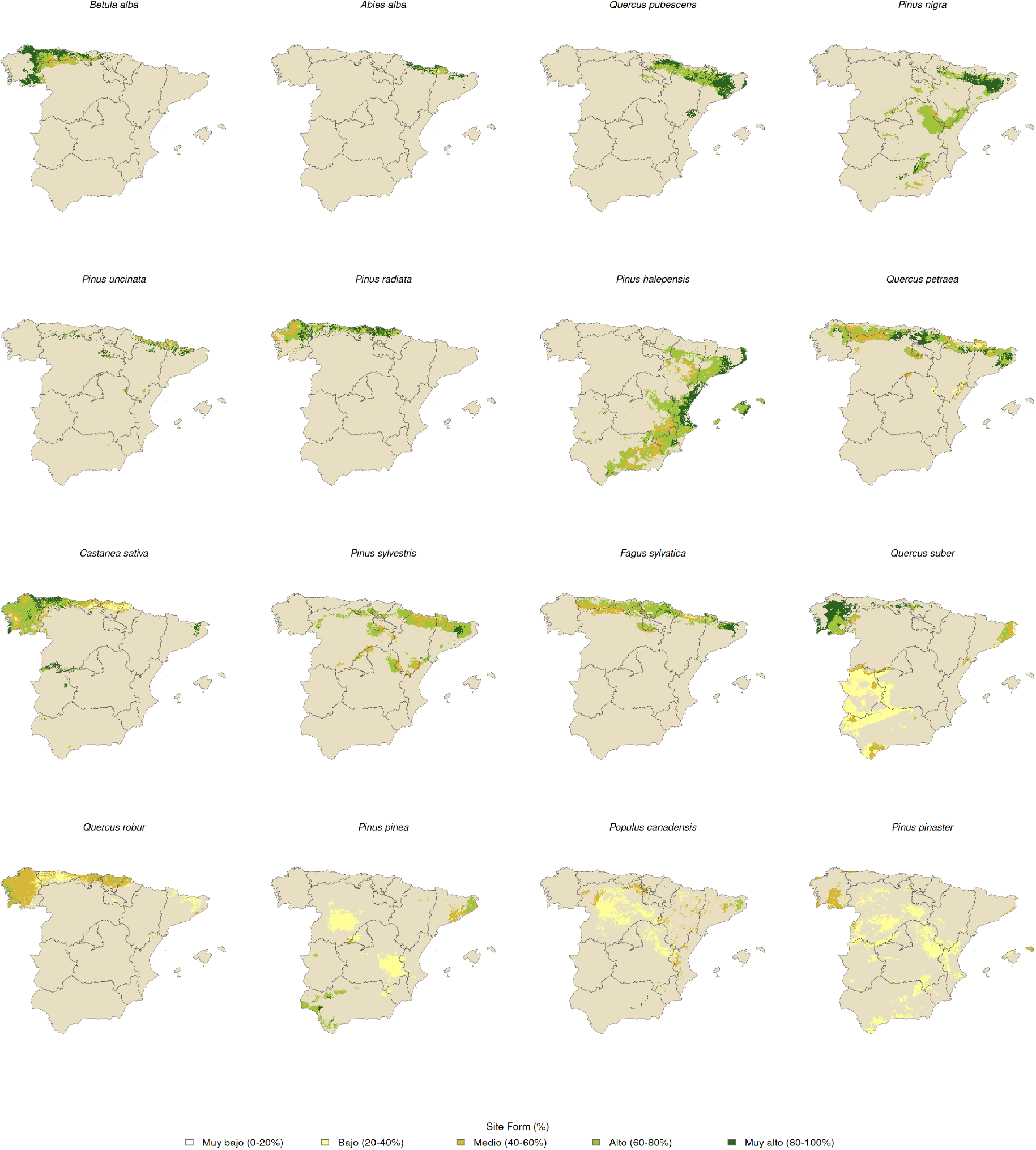
Reference-period (1985–2014) Site Form within each species’ SF-mapped area, for the 16 applicable species. All maps share the same 0–100% colour scale, from white (0–20%, lowest productivity) through pale yellow (20–40%) and ochre (40–60%) to light green (60–80%) and dark green (80–100%, highest productivity).

**Figure 4.**
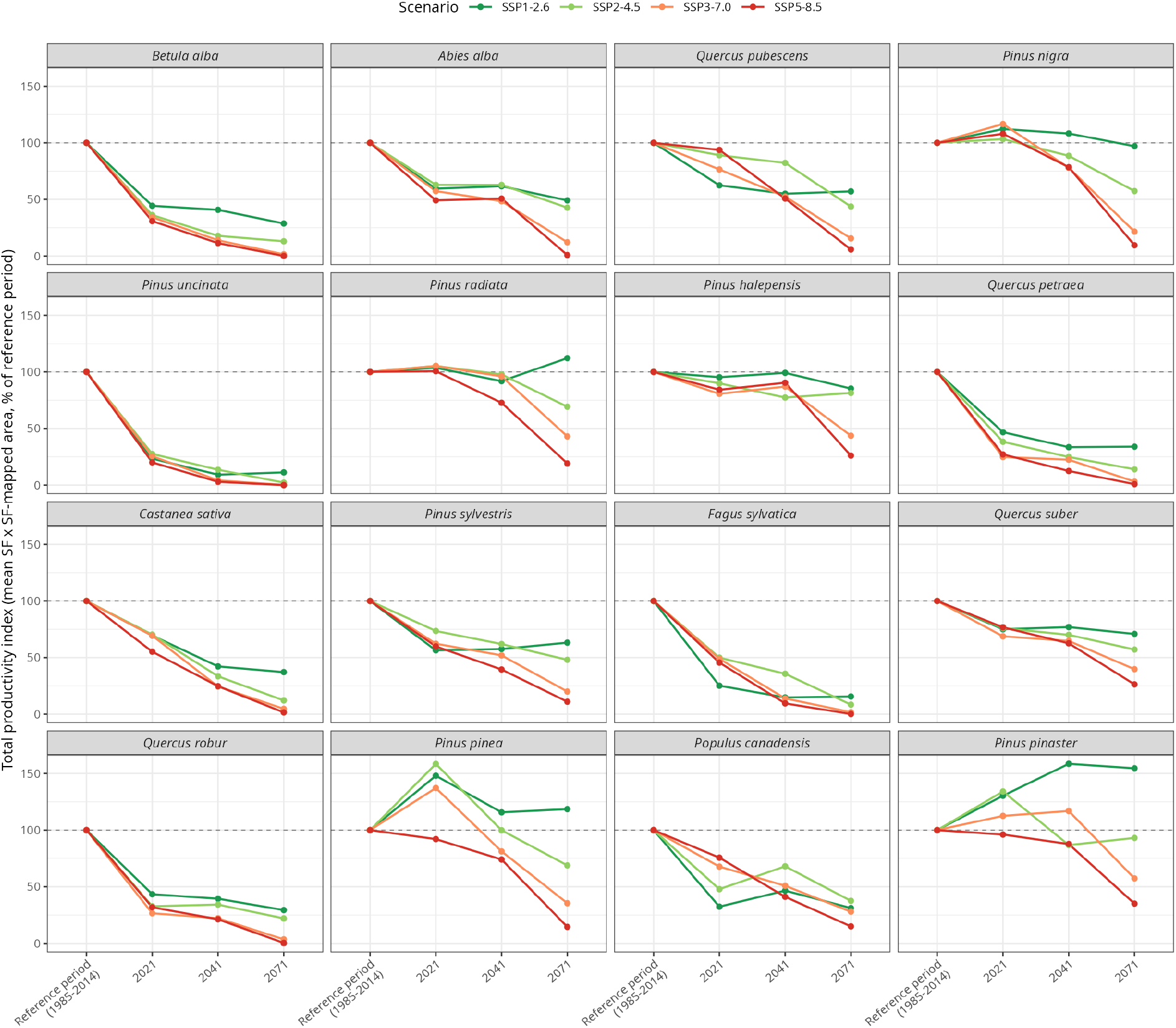
Total productivity index (mean SF *×* SF-mapped area, normalised to the reference-period baseline = 100%) for each of the 1 applicable species, across the three future horizons (2021–2050, 2041—2070, 2071—2100) and four emission scenarios (SSP1-2. to SSP-8.). Species are ordered from highest to lowest reference-period productivity (Table 2).

Fifteen of the 16 species lose total productivity under every scenario, with losses increasing progressively from SSP1-2.6 to SSP5-8.5 and, for most species, already substantial even under the lowest-emission pathway. Six species collapse to less than 1% of their reference-period value by 2071–2100 under SSP5-8.5: *Pinus uncinata* (a high-Pyrenean specialist, effectively eliminated), *Betula alba, Fagus sylvatica, Quercus robur, Quercus petraea* and *Abies alba* — all Euro-Siberian, montane, or sub-Atlantic species at the warm/dry margin of their European range. A second tier loses 85-98%: *Castanea sativa, Quercus pubescens, Pinus nigra, Pinus sylvestris, Pinus pinea* and *Populus canadensis*. A third, comparatively less-affected tier — *Pinus radiata, Pinus halepensis* and *Quercus suber* — still loses 74–81%. For four of these fifteen species (*Pinus nigra, Pinus pinea, Quercus pubescens, Populus canadensis*), the underlying SF-mapped area is stable or even growing (Section 4.3); the collapse in total productivity is driven entirely by mean SF within that area, a signal a simple range map would miss entirely. *Pinus pinaster* gains total productivity only under SSP1-2.6, reaching up to 175% of its reference-period value by 2071–2100; under the three higher-emission scenarios it instead loses productivity (93% under SSP2-4.5, 57% under SSP3-7.0, 35% under SSP5-8.5) — still the smallest loss of any of the 16 species under SSP5-8.5, but not the universal gain the SSP1-2.6 result alone would suggest. Constraining warming to SSP1-2.6 leaves total productivity comparatively intact for *Pinus nigra, Pinus pinea, Pinus radiata, Pinus halepensis* and *Quercus suber* (71–119% retained) but does little to prevent severe losses for the Euro-Siberian/montane group.

As an illustrative example, Figure 5 shows the spatial maps behind *Betula alba*’*s* collapse, currently the most productive species in the study (mean SF 66.7% over 59,943 mapped pixels along the humid Atlantic fringe of Galicia, Asturias and Cantabria). Reading the maps by colour alone can be misleading: the remaining pixels stay medium-to-dark green (60–100% SF) in every panel, giving the visual impression of sustained high productivity — indeed, mean SF among the survivors is unchanged or even slightly higher than the reference-period value. What actually collapses, and what Figure 4 s combined index quantifies directly, is the *extent* of the coloured area, which contracts from left to right (increasing emissions) and top to bottom (later horizons), down to a sliver of 37 pixels by 2071–2100 under SSP5–8.5 — a species that looks stable on a colour-only reading of its maps, and even in its own mean SF, but has in fact lost essentially all of its total productivity. Equivalent reference-period-to-future maps for all other applicable species are provided in Supplementary Figures S1–S17.

**Figure 5.**
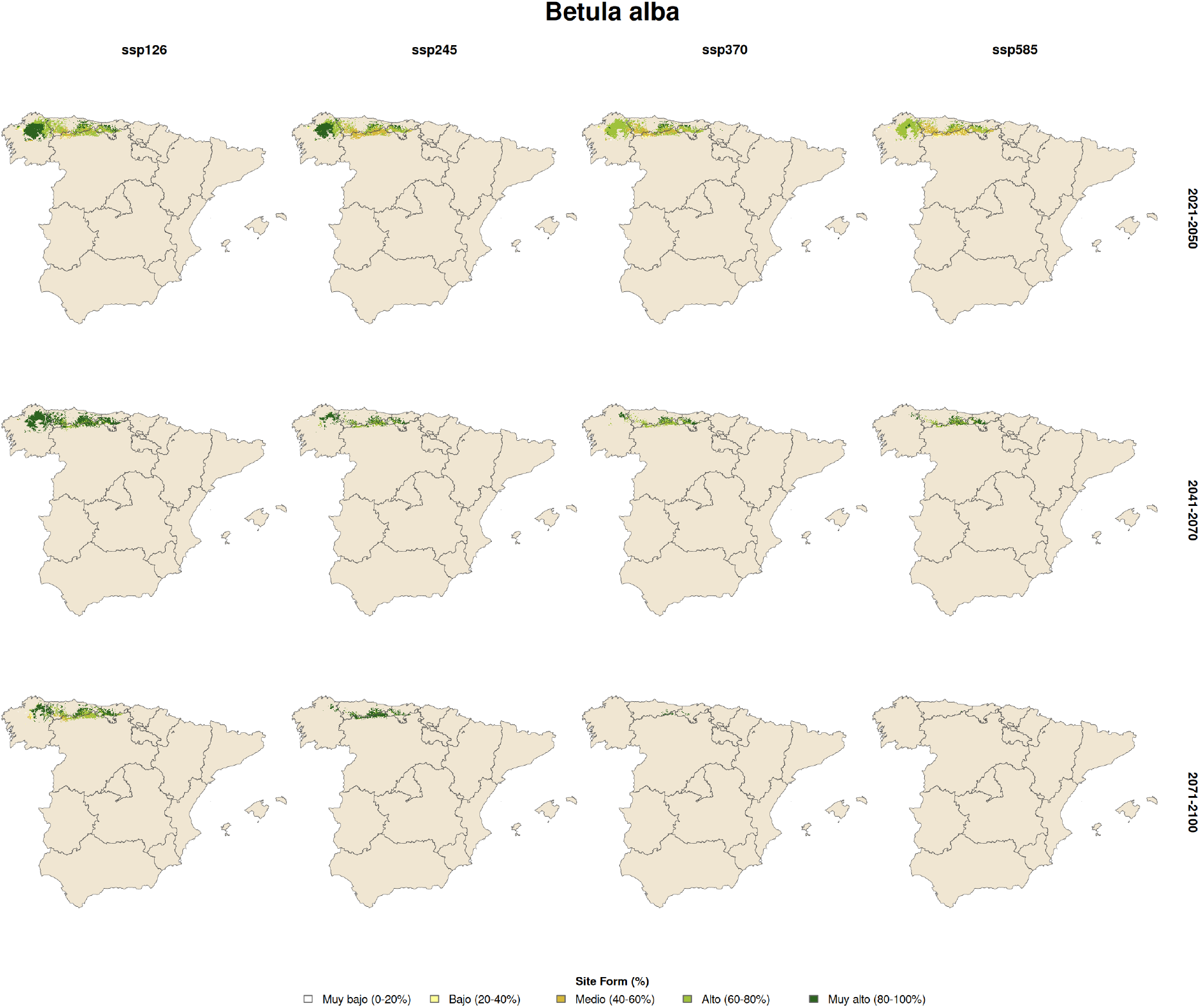
Ensemble-median (10 ESMs) Site Form (%) maps for Betula alba under four emission scenarios (columns) and three future horizons (rows). Colour encodes SF value only; the shrinking extent of the coloured area itself — not its colour, and not visible from mean SF alone — is what Figure 4’s combined index quantifies as a near-total collapse in total productivity (reference-period baseline: 59, 943 mapped pixels, mean SF 66.7%; SSP-8.5, 2071–2100: 37 pixels, mean SF 81.6%).

## 4. Discussion

### 4.1. Bioclimatic control of forest site productivity

The high validated skill reported above confirms that landscape-scale Site Form variation is largely bioclimatically controlled, consistent with prior work linking growth to temperature/precipitation gradients across Europe (Peñuelas et al., 2017; Ruiz-Benito et al., 2013). Lower performance for *Q. faginea, Q. ilex*, and *Q. pyrenaica* likely reflects their wide ecological amplitude and coppice-management history, which regional climate variables cannot capture (Calama et al., 2024). Sign patterns are ecologically informative: BIO14 precipitation of the driest month) is negative for drought-adapted species (*P. halepensis, J. thurifera*) — wetter sites are occupied by competing species that suppress site quality — but positive for temperate species (*B. alba, F. sylvatica*), reflecting genuine summer-drought limitation. The near-universal negative sign of BIO41/BIO42 (evapotranspiration; Section 3.1) points to atmospheric water demand penalising productivity independently of total precipitation, consistent with moisture stress as the dominant Mediterranean constraint. BIO13, despite being the single most frequently retained predictor overall, plays a similarly important but ecologically less clear-cut role: its weaker sign consistency suggests a more context-dependent relationship between wet-season surplus and growth, favourable for most species but occasionally offset by waterlogging or competitive release for drought-adapted specialists.

### 4.2. Model selection, ensemble averaging and methodological considerations

Neither AIC nor BIC consistently outperformed the other externally (7 vs. 6 species, MEAN best for 4), cautioning against a single information criterion as a universal rule. Where MEAN substantially outperformed both individual models (*P. halepensis, P. radiata, B. alba*), the two criteria evidently captured complementary signal (Burnham and Anderson, 2002); because its adoption was conditioned on outperforming both candidates in external validation rather than assumed a priori, MEAN is an empirically validated choice, not an unconditional averaging rule. The large AIC–BIC spread for some species (*P. pinea*: BIC 1.08 vs. AIC 7.84) suggests over-fitting under AIC or misspecification under BIC — an open question spatial cross-validation (Roberts et al., 2017) would help clarify.

SF conflates site quality with stand structure and management, introducing noise for intensively managed species (Calama et al., 2024), and bioclimatic multi-collinearity means coefficients should be read as predictive rather than causal. The habitat-suitability mask already gives an explicit, quantitative control on extrapolation, since a value is only reported where an SDM fitted independently of the SF regression supports the species’ presence under that climate. A continuous refinement — a per-pixel Eultivariate Environmental Similarity Surface [MESS; Elith et al. (2010)] or Mahalanobis distance — remains a natural next step to grade applicability *within* the SDM-suitable area, rather than to define its boundary.

### 4.3. Patterns of decline and resilience under future climate

Two distinct mechanisms underlie the near-total collapse in total productivity that Figure 4 shows for most species. *Pinus nigra* and *Pinus pinea* lose the great majority of their mean SF while their SF-mapped area barely changes: the footprint persists, but climate conditions inside it deteriorate almost to the calibration floor, and the combined index collapses accordingly. *Quercus pubescens* and *Populus canadensis* show the opposite mechanism: their SF-mapped area actually *grows* even as mean SF within it collapses, yet the index still falls to a small fraction of its reference-period value because the productivity term dominates the product. Because the habitat-suitability mask is fitted independently of the SF regression Section 2.5), this decomposition into area and productivity is a genuine, trackable signal rather than a modelling artefact, and a simple presence/absence range-shift map — or an area-only index — would have missed the productivity collapse for all four species.

For *Betula alba* and *Quercus robur*, decomposing the index resolves an apparent paradox: mean SF alone, computed over an ever-shrinking remnant, would have read as stable or mildly positive, and an area-only index would have flagged only these two species as catastrophic; it is the combined index in Figure 4 that correctly identifies both as near-total collapses while also capturing the four species above, whose area alone looked stable or improving. *Pinus radiata, Fagus sylvatica* and *Pinus uncinata* show a similar area-driven collapse, compounding already substantial mean-SF losses. *Pinus pinaster* is the one partial exception: its mean SF improves and its SF-mapped area contracts only moderately, so under SSP1-2.6 the gain outweighs the contraction and the combined index rises rather than falls; under the three higher-emission scenarios it still declines like every other species, but comparatively the mildest decline of the sixteen under SSP5-8.5.

### 4.4. Implications for carbon sequestration under climate change

SF is an indirect proxy for carbon-absorption capacity, not a flux measure: sequestration also depends on stand age, density, structure and management (Pan et al., 2011; Ruiz-Benito et al., 2013; Montero et al., 2005), and converting SF to absolute carbon requires species-specific allometric coupling — a priority future line, complicated by nonlinearities near distribution extremes, species-specific scaling, and competition in mixed stands.

The near-universal collapse in total productivity has direct accounting implications. For the six species collapsing to below 1% of their reference-period value (Section 3.3) — *P. uncinata, B. alba, F. sylvatica, Q. robur, Q. petraea and A. alba* — the carbon sink currently attributed to these forest types would be essentially eliminated under SSP5-8.5, whether the collapse is driven primarily by loss of mean SF, loss of SF-mapped area, or both. *P. nigra* and *P. pinea* illustrate a subtler risk: their SF-mapped area barely changes, so a static, area-based carbon inventory would keep crediting them at reference-period productivity even as their actual output collapses by over 90%. *Q. pubescens* and *P. canadensis* show the reverse framing risk: their growing range could be read as an expanding carbon asset, when in fact their total productivity within that range falls by 85-94%. *P. halepensis, P. radiata* and *Q. suber* retain the largest share of their reference-period productivity among the declining species (still losing 74– 81% by 2071–2100 under SSP5-8.5), but also carry increased wildfire risk under warmer, drier conditions. 0nly *P. pinaster* shows a genuine gain in carbon-relevant productivity, but only under the lowest-emission scenario; under higher-emission scenarios it declines like the rest, albeit less severely than any other species.

## 5. Conclusions

Projected under a high-emission future, the site productivity of Iberian forests faces a widespread, climate-driven collapse rather than a gradual decline: of the 16 species for which spatially explicit projections were possible, fifteen lose total productivity and six lose essentially all of it, with only one species gaining. We developed and validated species-specific bioclimatic regression models for Site Form across 21 Iberian tree species; 17 reached 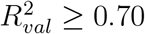 and their coefficients are used for spatial application. Bioclimatic variables explain 78% of landscape-scale SF variance on average (range 0.44–0.97). External validation against independent INIA data selected AIC for 7 species, BIC for 6, and MEAN for 4, reducing external RMSE by up to 59% relative to the best individual model where applicable. Assessed from each species’ actual final model rather than AIC alone, BIO13 (precipitation of the wettest month) is the most frequently retained predictor (15/17), followed by BIO04 (14/17) and BIO11 (13/17, predominantly negative); BIO10 and BIO18 were never retained. BIO44 is the most sign-consistent predictor among frequent variables (92% positive), while BIO41 and BIO42 are predominantly negative (89% and 91%); BIO14, the flagship predictor under AIC alone, is retained in only 11 of 17 final models with a near-even sign split, reflecting genuine ecological ambivalence rather than a universal role.

Final maps, for the reference period baseline and every one of 120 future scenario-period-ES combinations, rescale each prediction to the species’ calibration range and restrict it to the area an independently fitted species distribution model identifies as climatically suitable — an explicit, quantitative control on extrapolation risk at every pixel, rather than an implicit assumption that the regression remains valid everywhere. Applying this control to the spatial products themselves, rather than only to the statistical fit, also surfaced two cases — *Juniperus thurifera* and, more severely, *Pinus pinaster* under its AIC model — where an excellent 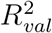 did not guarantee a usable spatial prediction; *J. thurifera* was excluded and *P. pinaster*’s reference-period and future projections were rebuilt from its less extreme) BIC model, a correction that would not have been visible from validation statistics alone.

Of the 16 species with usable reference-period maps, a combined index tracking both the extent and the mean productivity of each species’ SF-mapped area shows that fifteen lose total productivity under every scenario, six of them — *Betula alba, Quercus robur, Quercus petraea, Abies alba, Fagus sylvatica* and *Pinus uncinate —* collapsing to below 1% of their reference-period value by 2071–2100 under SSP5-8.5, and a further six — *Castanea sativa, Quercus pubescens, Pinus nigra, Pinus sylvestris, Pinus pinea* and *Populus canadensis* — losing 85–98%. Decomposing this index revealed two distinct decline mechanisms invisible to either component alone: productivity collapsing within an essentially stable range (*P. nigra, P. pinea*),and productivity collapsing even as the projected SF-mapped area expands (*Q. pubescens, P. canadensis*). A further two species (*B. alba, Q. robur*) show this same area-driven collapse from the opposite direction: an apparent mean-SF increase or mild decline that would look like resilience without also tracking area. 0nly *Pinus pinaster* gains total productivity, and only under SSP1–2.6 up to 175% of its reference-period value by 2071-2100); under higher-emission scenarios it declines like the rest of the sixteen species, though less severely. Constraining warming to SSP1-2.6 leaves total productivity comparatively intact for the Mediterranean pine/oak group (*P. nigra, P. pinea, P. radiata, P. halepensis, Q. suber*; 71–119% retained) but does little to prevent severe losses for the Euro-Siberian/montane group.

These validated, extrapolation-aware models and spatial products provide a direct route to dynamic, climate-aware carbon-uptake estimates for Iberian forests, integrable into the VisoCCarbono/OECC carbon calculator. Priority future research includes: coupling SF projections with species-specific allometric or growth models to translate percentile shifts into absolute carbon-flux estimates; quantifying the CO_2_ fertilisation effect; extending the framework to mixed stands; and applying spatial cross-validation and continuous MESS/Mahalanobis applicability surfaces within the SDM-suitable domain.

## Data availability

Site Form rasters are available from INIA and URJC upon reasonable request. All the research products are available at https://doi.org/10.5281/zenodo.21833757.

## Acknowledgements

We thank the Instituto Nacional de Investigación y Tecnología Agraria y Alimentaria INIA-CSIC) for providing the Site Form raster data derived from the Third Spanish National Forest Inventory and Eukene Azpitarte for growth data Populus × canadensis stands. This work was carried out within the VisoCCarbono project (“Plataforma de Visualización y Cálculo de Sumideros de Carbono Resilientes para la Optimización de la Mitigación del Cambio Climático en base a Escenarios Locales de Clima Futuro”, ref. CPP2022-009858), funded by the Spanish Agencia Estatal de Investigación inisterio de Ciencia, Innovación y Universidades) under the 2022 Public-Private Collaboration call of the Plan Estatal de Investigación Científica, Técnica y de Innovación 2021-2023, within the Plan de Recuperación, Transformación y Resiliencia, with support from the European Union — NextGenerationEU.

## Author contributions

**Marta Fernández-Pastor**^**⋆**^: Conceptualization, Data curation, Formal analysis, Writing — Original Draft, Writing — Review & Editing. **Gonzalo Rodríguez-Ruiz**^**⋆**^ : Conceptualization, Data curation, Formal analysis, Writing — Original Draft, Writing — Review & Editing. **Robert Monjo**: Methodology, Software, Formal analysis, Writing — Original Draft. **María del Carre**: Conceptualization, Project administration, Supervision. **Ana I. Hernández-Parada**: Conceptualization, Writing — Review & Editing. **Carlos Prado-López**: Data curation, Software (generation of past-climate and projected bioclimatic variable maps). **Raúl García Valdés**: Data curation, Resources (National Forest Inventory data used to model some species). **Darío Redolat**: Software, Writing — Review & Editing. **Eulogio Chacón**: Methodology (species distribution modelling), Writing — Review & Editing. **Jaime Ribalaygua**: Methodology (bioclimatic downscaling), Supervision.

^⋆^Marta Fernández-Pastor and Gonzalo Rodríguez-Ruiz contributed equally to this work.

## Appendix A. Bioclimatic predictor definitions and model performance

This appendix collects three reference tables kept out of the main narrative to keep it concise. Table A.3 defines the 25 bioclimatic predictor codes used throughout the analysis. Table A.4 gives the full stepwise-AIC regression performance (sample size, number of predictors, adjusted and validated R^2^, hold-out RMSE) for all 21 species, underlying the summary statistics reported in the main text. Table A.5 gives the corresponding external-validation RMSE for the three candidate approaches (BIC, AIC, MEAN) and the model selected for each of the 17 applicable species, complementing the specific examples discussed in the main text.

**Table A.3.** Description of the predictors (Bioclimatic variables) used in the analyses.

| Cod | Description |
| --- | --- |
| BIO01 | Annual Mean Temperature |
| BIO02 | Mean Diurnal Range (Mean of monthly (max temp - min temp)) |
| BIO03 | Isothermality (BIO2/BIO7) ($\times 100$) |
| BIO04 | Temperature Seasonality (standard deviation $\times 100$) |
| BIO05 | Max Temperature of Warmest Month |
| BIO06 | Min Temperature of Coldest Month |
| BIO07 | Temperature Annual Range (BIO5-BIO6) |
| BIO08 | Mean Temperature of Wettest Quarter |
| BIO09 | Mean Temperature of Driest Quarter |
| BIO10 | Mean Temperature of Warmest Quarter |
| BIO11 | Mean Temperature of Coldest Quarter |
| BIO12 | Annual Precipitation |
| BIO13 | Precipitation of Wettest Month |
| BIO14 | Precipitation of Driest Month |
| BIO15 | Precipitation Seasonality (Coefficient of Variation) |
| BIO16 | Precipitation of Wettest Quarter |
| BIO17 | Precipitation of Driest Quarter |
| BIO18 | Precipitation of Warmest Quarter |
| BIO19 | Precipitation of Coldest Quarter |
| BIO41 | Annual evapotranspiration |
| BIO42 | Evapotranspiration of the driest month |
| BIO43 | Evapotranspiration of the wettest or rainiest month |
| BIO44 | Evapotranspiration of the warmest month |
| BIO45 | Evapotranspiration of the coldest month |
| BIO46 | Seasonality of evapotranspiration ((Max - min) annual ) |

**Table A.4.** Stepwise AIC regression performance for 21 Iberian tree species ordered by 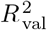.

| Species | $n$ | $K$ | $R^2_{\text{adj}}$ | $R^2_{\text{val}}$ | RMSE |
| --- | --- | --- | --- | --- | --- |
| **Pines** |  |  |  |  |  |
| *Pinus pinaster* | 5000 | 23 | 0.97 | 0.97 | 0.62 |
| *Pinus halepensis* | 5000 | 22 | 0.84 | 0.86 | 0.56 |
| *Pinus pinea* | 5000 | 22 | 0.86 | 0.86 | 0.69 |
| *Pinus radiata* | 5000 | 20 | 0.83 | 0.82 | 0.91 |
| *Pinus uncinata* | 3065 | 18 | 0.79 | 0.77 | 1.36 |
| *Pinus nigra* | 5000 | 22 | 0.72 | 0.72 | 1.31 |
| *Pinus sylvestris* | 5000 | 21 | 0.72 | 0.72 | 1.07 |
| **Oaks** |  |  |  |  |  |
| *Quercus pubescens* | 5000 | 21 | 0.90 | 0.91 | 0.84 |
| *Quercus robur* | 5000 | 22 | 0.88 | 0.87 | 0.60 |
| *Quercus suber* | 5000 | 16 | 0.79 | 0.77 | 0.44 |
| *Quercus petraea* | 4717 | 22 | 0.74 | 0.75 | 1.23 |
| *Quercus pyrenaica*<sup>†</sup> | 5000 | 21 | 0.64 | 0.63 | 0.89 |
| *Quercus ilex*<sup>†</sup> | 5000 | 20 | 0.62 | 0.59 | 0.32 |
| *Quercus faginea*<sup>†</sup> | 5000 | 19 | 0.41 | 0.46 | 1.10 |
| **Deciduous broadleaves** |  |  |  |  |  |
| *Betula alba* | 5000 | 22 | 0.93 | 0.93 | 0.81 |
| *Populus canadensis* | 2262 | 17 | 0.90 | 0.89 | 1.08 |
| *Fagus sylvatica* | 5000 | 21 | 0.84 | 0.83 | 0.92 |
| *Castanea sativa* | 5000 | 21 | 0.80 | 0.80 | 0.71 |
| *Populus nigra*<sup>†</sup> | 2595 | 17 | 0.64 | 0.67 | 1.30 |
| **Other conifers** |  |  |  |  |  |
| *Juniperus thurifera* | 5000 | 22 | 0.92 | 0.92 | 0.16 |
| *Abies alba* | 908 | 14 | 0.69 | 0.69 | 1.17 |

$n$ : pixels used for model fitting (max. 5,000). $K$ : selected predictors. $R^2_{\text{adj}}$ : adjusted $R^2$ on training set (80%). $R^2_{\text{val}}$ : $R^2$ on independent hold-out (20%). RMSE: root mean squared error on hold-out (SF units). †: $R^2_{\text{val}} < 0.70$ ; coefficients not applied for spatial projection. *Abies alba* ( $R^2_{\text{val}} = 0.69$ ) is retained as the value rounds to 0.70 at one decimal place.

**Table A.5.** External validation of the three candidate models against independent INIA forest data (Aguirre et al., 2022) for the 17 species with 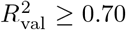.

| Species | RMSE<sub>BIC</sub> | RMSE<sub>AIC</sub> | RMSE<sub>mean</sub> | Selected |
| --- | --- | --- | --- | --- |
| **Pines** |  |  |  |  |
| *P. pinaster* | 4.72 | **3.87** | 4.18 | AIC |
| *P. halepensis* | 1.58 | 1.35 | **0.65** | MEAN |
| *P. pinea* | **1.08** | 7.84 | 3.62 | BIC |
| *P. radiata* | 3.75 | 4.14 | **0.99** | MEAN |
| *P. uncinata* | 4.23 | **1.59** | 2.76 | AIC |
| *P. nigra* | **1.76** | 2.43 | 2.06 | BIC |
| *P. sylvestris* | **1.16** | 1.96 | 1.46 | BIC |
| **Oaks** |  |  |  |  |
| *Q. petraea* | **1.42** | 3.29 | 2.00 | BIC |
| *Q. pubescens* | **1.86** | 2.97 | 2.37 | BIC |
| *Q. robur* | 3.76 | **1.48** | 1.49 | AIC |
| *Q. suber* | 1.56 | **1.14** | 1.35 | AIC |
| **Deciduous broadleaves** |  |  |  |  |
| *B. alba* | 5.20 | 7.87 | **1.65** | MEAN |
| *Pop. canadensis* | 3.96 | **1.95** | 2.91 | AIC |
| *F. sylvatica* | 2.73 | 3.98 | **1.22** | MEAN |
| *C. sativa* | 3.31 | **1.45** | 2.21 | AIC |
| **Other conifers** |  |  |  |  |
| *J. thurifera* | **3.61** | 4.12 | 3.86 | BIC |
| *A. alba* | 2.07 | **1.78** | 1.90 | AIC |

RMSE in SF units (m). The selected model (lowest external RMSE) is shown in bold.

